# Hyperactive Rac1 drives MAPK-independent proliferation in melanoma by assembly of a mechanosensitive dendritic actin network

**DOI:** 10.1101/326710

**Authors:** Ashwathi S. Mohan, Kevin M. Dean, Stacy Y. Kasitinon, Tadamoto Isogai, Vasanth Siruvallur Murali, Sangyoon J. Han, Philippe Roudot, Alex Groisman, Erik Welf, Gaudenz Danuser

## Abstract

Cancer cells use a variety of mechanisms to subvert growth regulation and overcome environmental challenges. Often, these same mechanisms enable cancer cells to also develop resistance to targeted therapies. Here, we describe how a hyperactivating mutation of the Rac1 GTPase (Rac1^P29S^) harnesses Rac1’s role as a regulator of actin polymer assembly to sustain cell cycle progression in growth limiting conditions. This proliferative advantage supports metastatic colonization of melanoma cells and confers insensitivity to inhibitors of the mitogen-activated protein kinase (MAPK) pathway, a frequent target for melanoma treatment. Rac1^P29S^ bypasses the MAPK axis through a mechanism that necessitates cell-matrix attachment, however, does not depend on integrin-mediated focal adhesion assembly and focal adhesion kinase signaling. Even without involvement of canonical adhesion signaling, cells carrying the Rac1^P29S^ mutation show elevated traction upon drug treatment and require mechanical resistance from their surrounding matrix to gain a proliferative advantage. We describe an alternative arm for cell mechanosensing, whereby actin polymerization against a matrix of minimal rigidity organizes biochemical cues to drive proliferative signals. Hyperactivation of Rac1 by the P29S mutation channels this pathway in melanoma through Arp 2/3-dependent formation of a constrained actin brush network that results in the inactivation of tumor suppressor NF2/Merlin. These data suggest an alternative mechanism for mechanosensitive growth regulation that can be hijacked by cancer cells to circumvent the adverse conditions of foreign microenvironments or drug treatment.

## Main

The RhoGTPase Rac1 is a central regulator of cellular functions like cytoskeletal organization and cell proliferation that underlie cancer development and progression ^1, 2^. The dependence on Rac1 for Ras-driven tumorigenesis and invasion ^3–6^ along with widely observed overexpression of Rac1 in many cancers ^7–11^ were early indicators that Rac1 is an essential factor in cancer progression. However, increasing evidence of deregulation of Rac1 activity suggests that Rac1 may not only be a necessary ingredient in cancer progression, but may function as a cancer driver itself. Guanine nucleotide exchange factors (GEFs) that activate Rac1 are frequently overexpressed or mutated in cancer ^2, 12^. For the set of 469 patient cases listed in The Cancer Genome Atlas’ Skin Cutaneous Melanoma database, a diverse array of mutations in the Rac1 GEFs, TIAM-1,2, PREX-1,2, Vav-1, and Dock 3, cumulatively occur at a frequency of 89.5%. Although less frequent, activating mutations in Rac1 itself have been found by recent genomic analyses in up to 9% of patients ^13, 14^. Of these, Rac1^P29S^, enriched in melanoma, is the most common ^15^. As a physiologically relevant, easily tractable perturbation of Rac1 activity, Rac1^P29S^ presents a valuable tool for better understanding the critical pathobiology of Rac1 hyperactivation in cancer progression.

Using a mouse model previously shown to reflect metastatic potential in melanoma patients ^16^, we found that subcutaneous xenograft tumors formed from A375 melanoma cells exogenously expressing Rac1^P29S^ (+P29S), Rac1^WT^ (+WT), and empty vector (+EV) showed no growth difference in the primary tumors (Fig. 1a), suggesting that survival and proliferation in the primary injection site are not affected by the Rac1^P29S^ mutation. Interestingly, although mice harboring these xenograft tumors exhibit metastases in various organs regardless of Rac1 mutation status or expression level, bioluminescence imaging (BLI) (Fig. 1b, c and Supplementary Fig. 1) and gross observation of metastatic lung nodules (Fig. 1d and Supplementary Fig. 2) revealed that the overall metastatic burden was significantly increased for mice with xenograft tumors expressing Rac1^P29S^. This suggests that the Rac1^P29S^ mutation confers a metastasis-specific growth advantage *in vivo*. Since one of the factors that must be overcome for metastasis is proliferation-limiting conditions at the site of colonization ^17^, we reasoned that expression of Rac1^P29S^ might confer a proliferative advantage to melanoma cells that have metastasized. We find that metastatic nodules in mice with Rac1^P29S^ tumors indeed have increased proliferation (Fig. 1e, f), suggesting that Rac1^P29S^ confers a proliferative advantage to melanoma cells in secondary sites that is not at play in the primary tumor.

**Figure 1.**
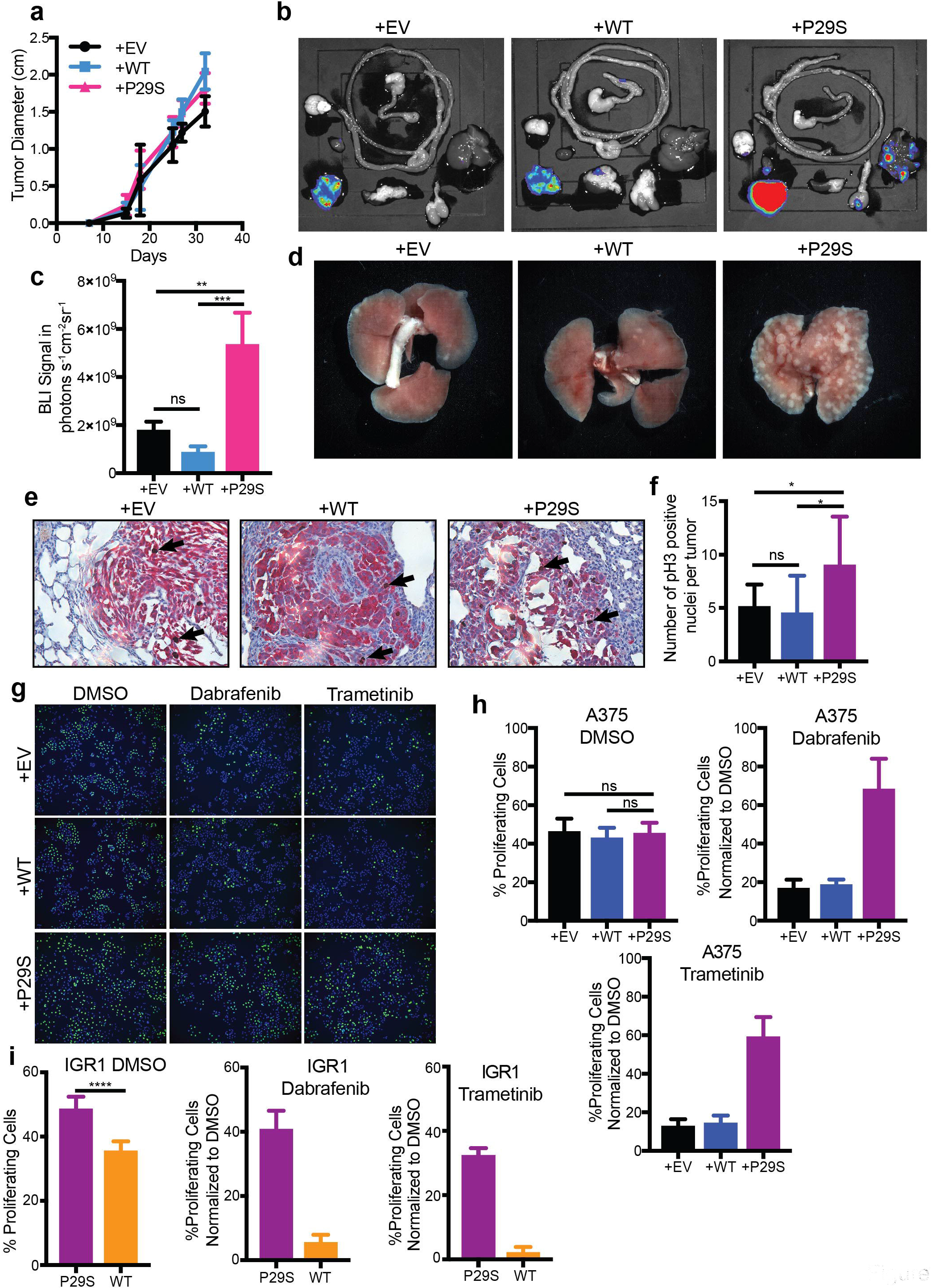
Rac1^P29S^ confers growth advantage to melanoma cells in metastases and upon MAPKi treatment. **a,** Growth of xenograft tumors in NSG mice from A375 melanoma cells exogenously expressing Rac1^P29S^ (+P29S), Rac1^WT^ (+WT), or empty vector (+EV) (n=20 mice per condition from two independent experiments). **b-d,** Endpoint metastatic burden in mice with tumors that were between 2.0-2.5 cm. (**b, c**) Bioluminescence imaging (BLI) signal detected in dissected organs. (**d**) Macrometastases in gross lungs post fixation in 10% neutral-buffered formalin for 48hrs (n=20 mice per condition from two independent experiments, see also Supplementary Fig. 1,2). **e,** Paraffin-embedded lungs were sectioned and dual-labeled with anti-dsRed antibody (red) for tumor nodule detection and anti-phospho Histone 3 for detection of mitotic nuclei (brown, representative examples indicated with arrows). **f,** All metastatic lung nodules controlled for size using Ilastik pixel classification were randomized and hand counted for phospho-Histone 3 positive nuclei to determine number of proliferating cells (n ≥ 9 metastatic lung nodules for each condition). **g, h,** Incorporation of fluorescently-labeled Edu into S-phase nuclei of A375 cells to detect proliferating cells (green) and Hoechst-labeled total nuclei (blue) to determine % proliferating cells upon 48 hr treatment with DMSO (0.003%), Dabrafenib (33.3 nM), and Trametinib (3.3 nM). **i,** Proliferation assay applied to IGR1 cells endogenously expressing Rac1^WT^ or Rac1^P29S^ upon 48 hr treatment with DMSO (0.1%), Dabrafenib (10,000 nM), and Trametinib (6.6 nM). For all Edu incorporation assays n=15 images were counted for each condition from three independent experiments. All data represent mean ± s.d. Statistical significance was assessed by one-way ANOVA with Tukey’ s multiple comparisons test (**c, f**) or a two-sample equal variance Student’ s *t*-test (**h, i**). *P<0.05, **P<0.01, ***P<0.001, ****P<0.0001.

To identify additional conditions under which the Rac1^P29S^ mutation confers a proliferative advantage, we tested if a proliferative mechanism might account for the drug resistance to BRAF and MEK inhibitors (MAPKi) recently observed in cells expressing Rac1^P29S 18^. Consistent with the comparable primary tumor growth rates observed *in vivo*, cells expressing Rac1^WT^ and Rac1^P29S^ exhibited comparable proliferation without treatment *in vitro* (Fig. 1g, h). Upon MAPKi treatment however, control cells suffered a massive drop in proliferation while cells expressing Rac1^P29S^ exhibited resistance to the proliferation inhibition (Fig. 1g, h, Supplementary Fig. 3). We also measured proliferation in response to MAPKi-treatment in IGR1 melanoma cells that carry the Rac1^P29S^ mutation endogenously (IGR1^P29S^) and IGR1 melanoma cells in which we reverted the native Rac1^P29S^ mutation to Rac1^WT^ using CRISPR-Cas9 genome editing (IGR1^WT^) (Supplementary Fig. 4). While IGR1^P29S^ cells proliferated at a slightly higher rate than IGR1^WT^ cells, the most prominent proliferative advantage for IGR1^P29S^ was noted upon MAPKi treatment (Fig. 1i). IGR1^P29S^ cells exhibited sustained proliferation under both BRAF and MEK inhibition, whereas IGR1^WT^ cells were no longer resistant (Fig. 1i), demonstrating that resistance to proliferation inhibition is driven by Rac1^P29S^. These data, along with our *in vivo* results, suggest that Rac1 hyperactivity creates a protective mechanism whereby melanoma cells continue to proliferate in spite of environmental or chemical conditions that otherwise reduce proliferation.

As proposed in previous studies ^14, 18^, we find that exogenous expression of Rac1^P29S^ results in slightly elevated MAPK signaling (Fig. 2a), suggesting that Rac1^P29S^ maintains cell proliferation by sustaining MAPK signaling following treatment with BRAF and MEK inhibitors. However, we find that phospho-MEK and phospho-ERK levels are suppressed in Rac1^P29S^ cells following MAPKi treatment (Fig. 2a), suggesting that signaling through the MAPK pathway remains sensitive to BRAF and MEK inhibition in Rac1^P29S^ cells even though proliferation is sustained. Additionally, we found that expression of the cell cycle checkpoint protein cyclin D1 was sustained upon suppression of MEK and ERK activity in cells expressing Rac1^P29S^, whereas cyclin D1 levels were highly sensitive to MEK and ERK suppression in control cells (Fig. 2a). This suggests that elevated MAPK signaling in Rac1^P29S^ cells might be de-coupled from Rac1^P29S’^s proliferative advantage. Indeed, even though direct inhibition of Erk activity with SCH772984 completely ablates ERK phosphorylation in Rac1^P29S^ cells just as in control cells (Fig. 2b), we find that cyclin D1 levels remain elevated and proliferation is sustained in Rac1^P29S^ cells (Fig. 2b,c). Thus we conclude that sustained proliferation - and consequently drug resistance - in cells expressing Rac1^P29S^ does not require MAPK or ERK signaling (Fig. 2d).

**Figure 2.**
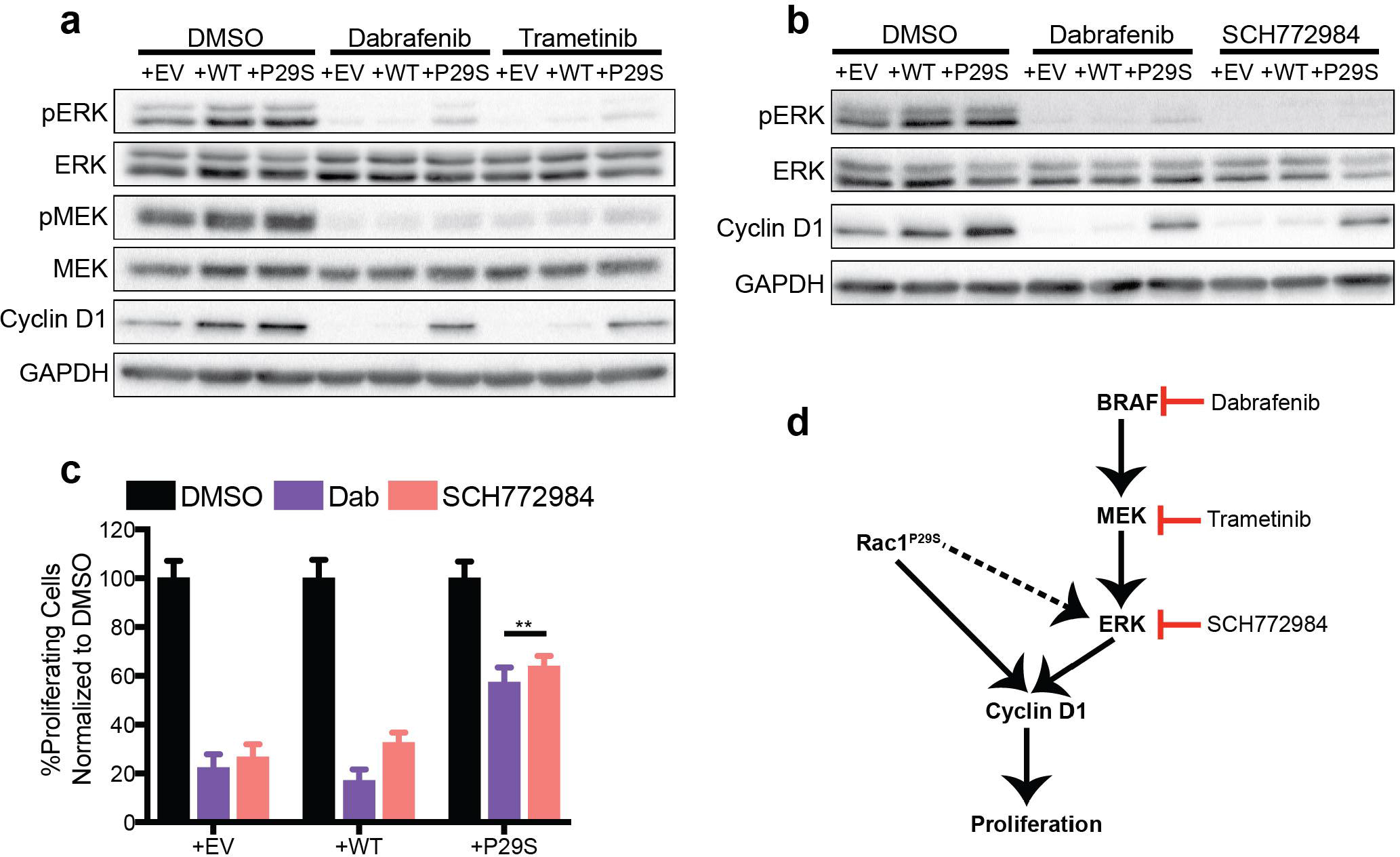
Rac1^P29S^ drives proliferation through a MAPK-independent pathway. **a, b,** MAPK signaling represented by phospho-ERK and phospho-MEK and cyclin D1 levels in A375 melanoma cells exogenously expressing Rac1^P29S^ (+P29S), Rac1^WT^ (+WT), or empty vector (+EV) upon 48 hr treatment with (**a**) DMSO (0.003%), Dabrafenib (33.3 nM), and Trametinib (3.3 nM) and (**b**) DMSO (0.003%), Dabrafenib (33.3 nM), and SCH772984 (100 nM). Data are representative of two independent experiments. **c,** Proliferation assay applied to A375 melanoma cells upon 48 hr treatment with DMSO (0.003%), Dabrafenib (33.3 nM), and SCH772984 (100 nM). For all Edu incorporation assays n=15 images were counted for each condition from three independent experiments. Statistical significance was assessed using a two-sample equal variance Student’ s *t*-test. Bar graphs represent mean ± s.d. **P<0.01. **d,** While Rac1^P29S^ increases MAPK signaling, its ability to drive the cell cycle and sustain proliferation are independent of the MAPK pathway and ERK activity.

Since the proliferative advantage imparted by Rac1^P29S^ is MAPK-independent, we sought alternative mechanisms that could support the robust proliferation observed in cells expressing Rac1^P29S^ (Supplementary Fig. 5a, b, c) ^19^. Given Rac1’s role in cytoskeletal remodeling, and the importance of cell-matrix adhesions in cancer cell proliferation ^20, 21^, we decided to investigate if Rac1^P29S^ altered focal adhesion formation and turnover. We observed that melanoma cell lines expressing endogenous Rac1^P29S^ had strikingly uncharacteristic numbers, distributions, and sizes of adhesions compared to cell lines expressing endogenous Rac1^WT^ (Fig. 3a). Introducing Rac1^P29S^ into cells via stable exogenous expression resulted in the formation of large adhesions (Extended Data Fig. 1a, b) that increased in size upon MAPKi treatment (Extended Data Fig. 1c, d). Thus, we hypothesized that Rac1^P29S^ overcomes growth suppression by altering cell-matrix adhesions and signaling. We find that the melanoma cells we evaluated are resistant to anoikis and proliferate like adherent cells in suspension (Extended Data Fig. 1e). However, unlike in adherent conditions, we find that cells expressing Rac1^P29S^ and control cells are equally sensitive to MAPKi treatment in suspension (Fig. 3b, Supplementary Fig. 5d, e). This suggests that the proliferative advantage conferred by Rac1^P29S^ is dependent on cell-matrix adhesion.

**Figure 3.**
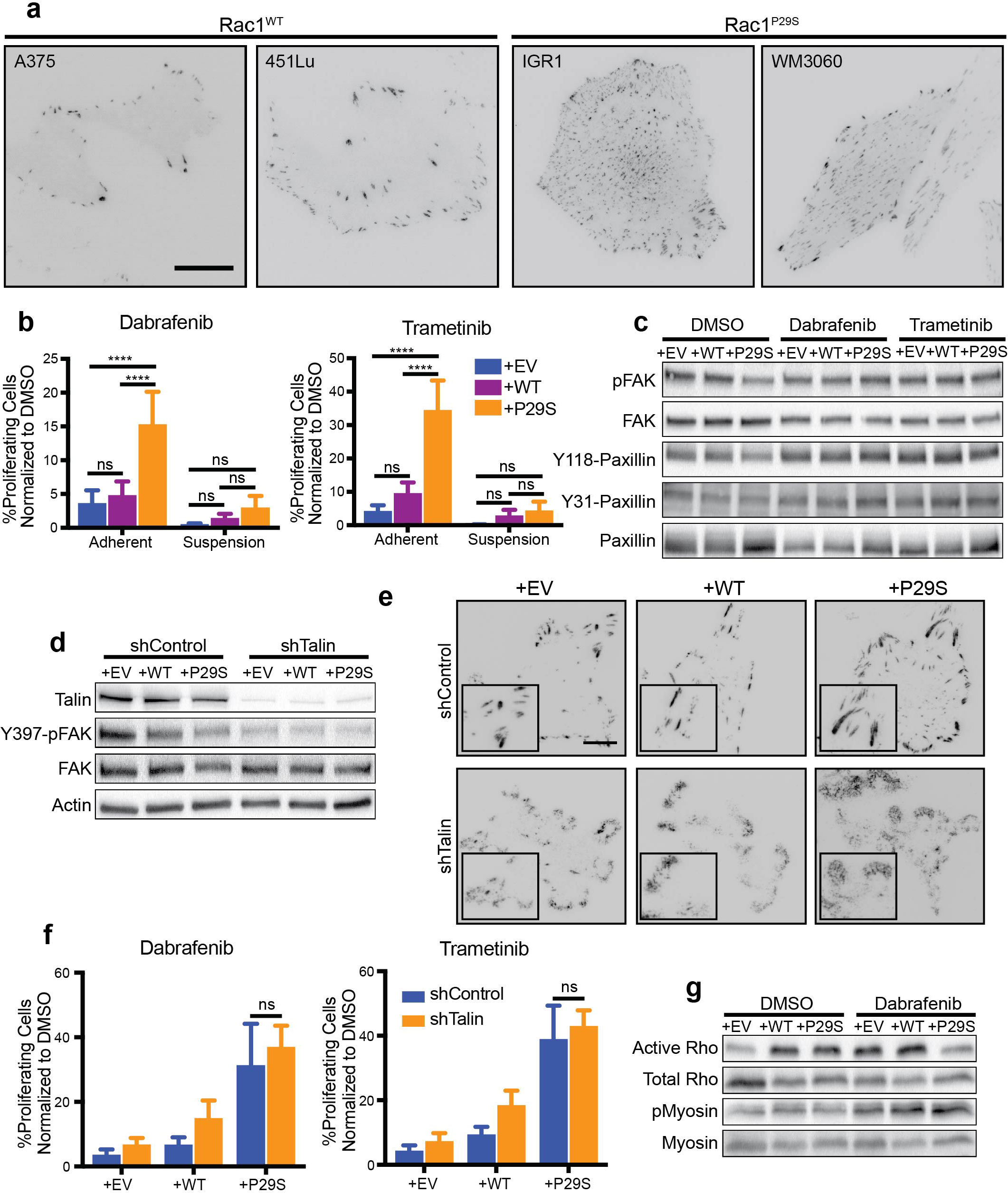
Rac1^P29S^ driven proliferation depends on cell-matrix attachment, but is independent of canonical focal adhesion signaling. **a,** Comparison of adhesion structures (inverted mNeonGreen-Paxillin signal) between cells endogenously expressing Rac1^WT^ versus Rac1^P29S^. Images are representative of the predominant phenotype observed across all cells for n=15 images acquired. Scale bar = 20 um. **b,** Proliferation assay applied to A375 melanoma cells exogenously expressing Rac1^P29S^ (+P29S), Rac1^WT^ (+WT), or empty vector (+EV) upon 48 hr treatment with Dabrafenib (33.3 nM) and Trametinib (3.3 nM). Cells were grown on tissue culture coated plastic to facilitate adhesion or in low-adhesion dishes to inhibit adhesion. Statistical significance was assessed using a 2-way ANOVA. ****P<0.0001. **c,** Focal adhesion kinase (FAK) and Paxillin signaling activities in A375 melanoma upon 48 hr treatment with DMSO (0.003%), Dabrafenib (33.3 nM), and Trametinib (3.3 nM) treatment. Data are representative of two independent experiments. **d, e** Disruption of focal adhesion complexes by stable expression of pGIPZ-TLN-V2LHS-56643 (shTalin) to achieve talin knockdown or pGIPZ-control-V2LHS (shControl). (**d**) Reduction of talin expression and FAK activity upon shTalin and shControl expression across all A375 cells lines expressing Rac1P29S (+P29S), Rac1^WT^ (+WT), or empty vector (+EV). (**e**) Immunofluorescence detection of focal adhesion disruption following expression of shTalin or shControl using anti-paxillin antibody in A375 cells with Rac1P29S (+P29S), Rac1^WT^ (+WT), or empty vector (+EV). Scale bar = 10 um. **f,** Proliferation assay applied to respective A375 cells expressing shControl or shTalin upon 48 hr treatment with Dabrafenib (33.3 nM) and Trametinib (3.3 nM). A two-sample equal variance Student’ s *t*-test was performed to compare proliferation between Rac1^P29S^ cells expressing shControl versus shTalin when treated with Dabrafenib (p=0.146) and Trametinib (p=0.184). **g,** Activation of RhoA (assessed by GST-RBD pull down) and myosin (assessed by phosphorylation of myosin light chain) in A375 cells expressing Rac1^P29S^ (+P29S), Rac1^WT^ (+WT), or empty vector (+EV) upon 48 treatment with DMSO (0.003%) and Dabrafenib (33.3 nM). Data are representative of two independent experiments. All bar graphs represent mean ± s.d.

To further test the dependence of Rac1^P29S^ proliferative signaling on cell-matrix adhesions, we evaluated cell-matrix signaling events. Curiously, signaling of focal adhesions via FAK and Paxillin activity did not appear noticeably different for cells expressing Rac1^P29S^ in the presence or absence of MAPKi treatment (Fig. 3c). To investigate this further, we disrupted focal adhesion formation by stable knockdown of Talin (Fig. 3d, e). Talin is an adhesion scaffolding protein that serves as the main molecular linker between integrins and the contractile actomyosin network that elicits adhesion maturation and signaling ^22^. As expected, Talin-knockdown resulted in reduced activation of the canonical adhesion-mediated signaling pathway characterized by phosphorylation of focal adhesion kinase (FAK; Fig. 3d). Strikingly, however, we found that the proliferative advantage conferred by Rac1^P29S^ was completely unperturbed by the knockdown of Talin (Fig. 3f, Extended Data Fig. 1f). This finding suggests that while cell attachment is necessary for the sustained proliferation observed in Rac1^P29S^ cells upon MAPKi treatment, Rac1^P29S^ drives proliferation through a pathway independent of canonical adhesion signaling. Consistent with a non-canonical mechanism of matrix engagement, we find that Rac1^P29S^ cells have reduced RhoA activity upon drug treatment (Fig. 3g). This is counter to the well-characterized focal adhesion signaling mechanism in which cellular contractility is increased, thereby facilitating heightened adhesion signaling ^20^. Likewise, while MAPKi treatment led to an expected overall elevation in myosin phosphorylation ^23^, this effect was not unique to cells expressing Rac1^P29S^ (Fig. 3g). Thus, Rac1^P29S^ mediates attachment-dependent proliferation through a mechanism independent of focal adhesions and actomyosin contractility (Supplementary Fig. 6).

Although the proliferative advantage in MAPKi-treated Rac1^P29S^ cells appears independent of actomyosin contractility, we observe a dependence of the proliferative advantage on matrix rigidity (Fig. 4a, Extended Data Fig. 2a, b). We cultured cells in pure collagen gels (~200 Pa) and collagen gels that were treated with ribose to increase collagen crosslinking and gel stiffness (~500 Pa) while collagen concentration was kept constant ^24^. We found that the proliferative advantage under MAPKi treatment is restored in Rac1^P29S^ cells in a stiffness-dependent manner while control cells continue to exhibit proliferation rates similar to those in suspension, regardless of the stiffness of the collagen gel (Fig. 4a, Extended Data Fig. 2a, b). Upon measuring the force cells exert on their matrix directly ^25^, we find that cells expressing Rac1^P29S^ have a pronounced increase in strain energy upon MAPKi treatment (Fig. 4b, Extended Data Fig. 2c). This is remarkable since upon MAPKi treatment, Rho activity is reduced and actomyosin contractility is not uniquely increased in Rac1^P29S^ cells (discussed above, Fig. 3g). Additionally, we note that mechanosensitive proliferation in MAPKi-treated Rac1^P29S^ cells spans a collagen rigidity range over a few hundred Pascals ^24^, whereas modulation of canonical mechanotransduction pathways spans a range an order of magnitude higher ^20^ (Supplementary Fig. 7). Thus, cells expressing Rac1^P29S^ engage mechanical cues from their environment upon MAPKi treatment to drive proliferation using a mechanosensitive pathway that is distinct from the well-established integrin-adhesion-actomyosin axis ^20,21^.

**Figure 4.**
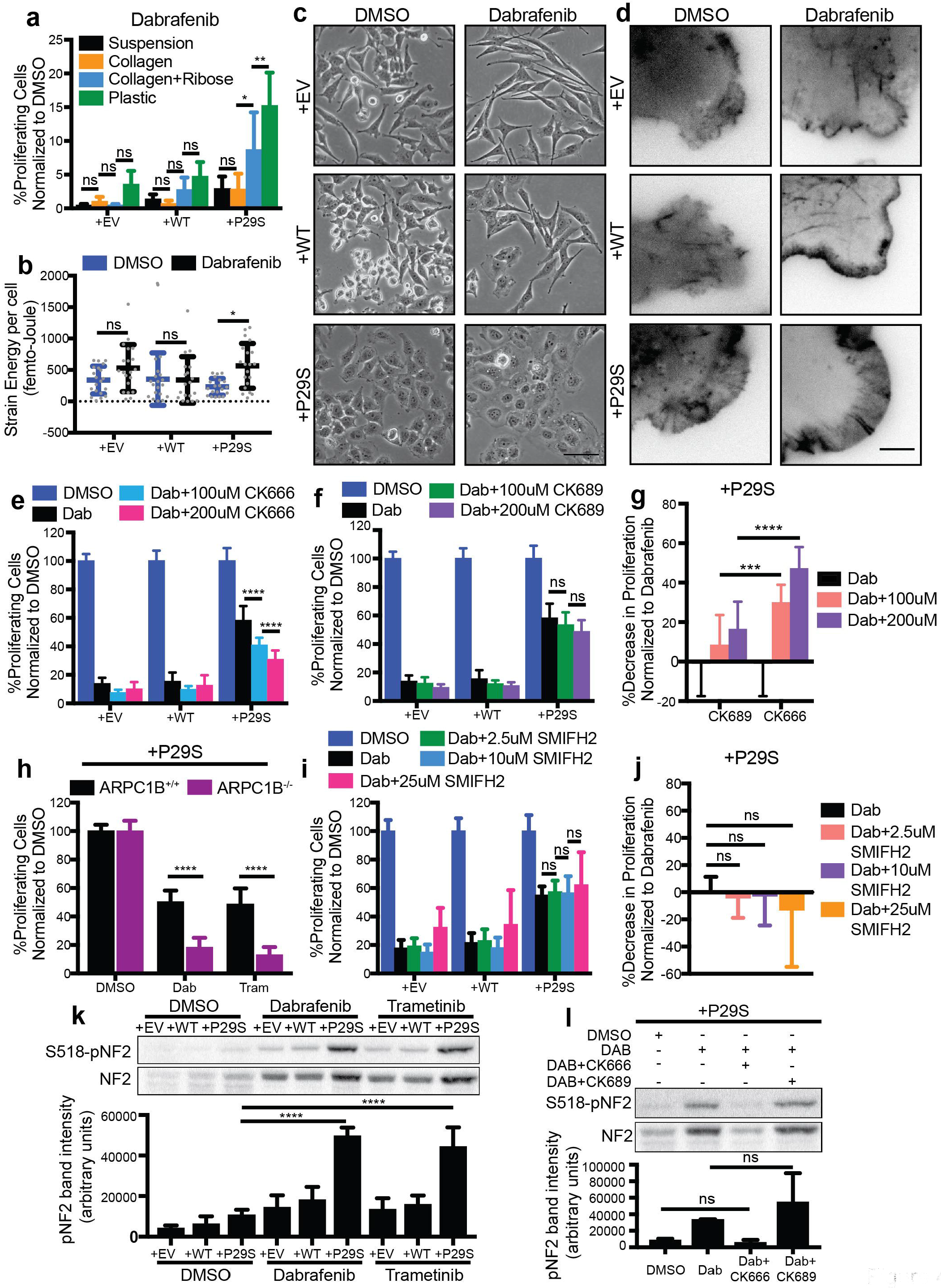
Rac1^P29S’^ s proliferative advantage is driven by enhanced assembly of a dendritic actin network, which inactivates the tumor suppressor NF2. **a,** Assessed by the proliferation assay, A375 cells expressing Rac1^P29S^ (+P29S) but not control cells expressing empty vector (+EV) or Rac1^WT^ (+WT) are resistant to proliferative suppression by MAPKi treatment in a stiffness-dependent fashion. Cell lines were cultured using indicated substrates and treated with Dabrafenib (33.3 nM) for 48hrs. **b,** Strain energy per cell determined by Traction Force Microscopy (TFM) increases upon MAPKi treatment for cells expressing Rac1^P29S^ (+P29S), but not control cells expressing empty vector (+EV) or Rac1^WT^ (+WT). Prior to imaging, cells were cultured on TFM substrates for 48 hrs with DMSO (0.003%) or Dabrafenib (33.3 nM) treatment. **c,** Cell morphology changes upon 48 hr Dabrafenib (33.3 nM) treatment (20x magnification). Control cell lines expressing empty vector (+EV) or Rac1^WT^ (+WT) adopt a spindle-like morphology, while cells expressing Rac1^P29S^ (+P29S) display enhanced spreading. Scale bar = 50 um. **d,** Changes in actin cytoskeletal architecture at cell edges following 48 hr Dabrafenib (33.3 nM) treatment. Contrast-inverted TIRF microscopy images (60x magnification) of mNeonGreen-Actin-C-35 construct stably co-expressed in A375 melanoma cells with Rac1^P29S^ (+P29S), Rac1^WT^ (+WT), or empty vector (+EV). Scale bar = 5 um. **e-h,** Proliferation assay applied to A375 melanoma cells expressing Rac1^P29S^ (+P29S), Rac1^WT^ (+WT), or empty vector (+EV) upon 48hr treatment with Dabrafenib (33.3 nM) along with indicated concentrations of (**e**) the Arp 2/3 complex inhibitor CK666 and (**f**) the control peptide CK689. (**g**) Decrease in proliferation relative to proliferation under Dabrafenib treatment alone. (**h**) Proliferation assay applied to A375 cells expressing Rac1^P29S^ (+P29S) with CRISPR/Cas9-mediated knock-out of the Arp2/3 regulatory component ARPC1B (ARPC1B^−/−^) and with control levels of ARPC1B (ARPC1B^+/+^). Both cell types were treated for 48 hrs with DMSO (0.003%), Dabrafenib (33.3 nM), and Trametinib (3.3 nM). **i, j,** Proliferation assay applied to A375 melanoma cells expressing Rac1^P29S^ (+P29S), Rac1^WT^ (+WT), or empty vector (+EV) upon 48 hr treatment with Dabrafenib (33.3 nM) along with indicated concentrations of the pan-formin inhibitor SMIFH2 (**i**). Absence of change in proliferation relative to proliferation under Dabrafenib treatment alone (**j**). **k, l** Total NF2 and inactive S518-phospho-NF2 in A375 melanoma cells exogenously expressing Rac1^P29S^ (+P29S), Rac1^WT^ (+WT), or empty vector (+EV) upon 48 hr treatment with (**k**) DMSO (0.003%), Dabrafenib (33.3 nM), and Trametinib (3.3 nM) and (**l**) dual Dabrafenib (33.3 nM) and CK666 (200 uM) or CK689 (200 uM) treatment. Data are representative of three independent experiments. Quantification by densitometry is the pixel area under the curve of band intensities. Statistical significance was determined using a two-way ANOVA and Tukey’ s multiple comparisons test for (**a, e-g, i, j**). A Kruskal-Wallis and Dunn’ s multiple-comparisons test was used to determine statistical significance for (**b**) due to non-normal distribution of the data determined using D’Agostino-Pearson omnibus normality test. An unpaired Student’ s t-test was used to determine statistical significance for (**h**). A one-way ANOVA and Tukey’ s multiple comparisons test were used in (**k and l**). The p-value in (**l**) for DMSO versus Dab+CK666 is 0.9970 and for Dab versus Dab+CK689 is 0.5049. All bar graphs represent mean ± s.d. *P<0.05, **P<0.01, ***P<0.001, ****P<0.0001.

We reasoned that contractility-independent mechanosensitivity and increases in traction strain observed in cells expressing Rac1^P29S^ upon MAPKi treatment could be caused by elevated actin polymerization ^26^, which might in turn regulate proliferation. We have previously shown that short-term inhibition of ERK abrogates lamellipodial actin polymerization ^27^. Consistently, we find that upon long-term MAPKi treatment, control cells become elongated spindles whereas cells expressing Rac1^P29S^ maintain and even enhance their spread morphologies (Fig. 4c). Upon closer inspection, we find that this enhanced cell spreading observed in Rac1^P29S^ cells upon MAPKi treatment is characterized by the development of wide, persistent lamellipodia (Fig. 4d, Movie S1). This change in cellular architecture, unique to Rac1^P29S^ cells upon drug treatment mirrors the Rac1^P29S^-specific increase in cell strain energy observed upon MAPKi treatment (Fig. 4b) and is consistent with the proliferative advantage unique to Rac1^P29S^ cells under drug treatment. Thus, we hypothesized that upon ERK inhibition, hyperactive Rac1^P29S^ sustains cell proliferation by enhancing dendritic network assembly in lamellipodia. We disrupted lamellipodia formation by inhibition of the Arp 2/3 complex, the nucleator of branched actin polymerization^28^. Indeed, pharmacological inhibition of Arp2/3 activity with compound CK666 reduces the proliferative advantage of Rac1^P29S^ cells upon MAPKi treatment, while treatment with the control peptide CK689 does not (Fig. 4e-g, Extended Data Fig. 2d, e). Additionally, abrogation of Arp 2/3 activity with CRISPR/Cas9-mediated genomic knockout of the ARPC1B regulatory domain ^29^ (Supplementary Fig. 8) also renders Rac1^P29S^ cells sensitive to MAPKi treatment (Fig. 4h, Extended Data Fig. 2f). Thus, we find that Arp2/3 activity is necessary to mediate the proliferative advantage conferred by Rac1^P29S^.

Upon inhibition of linear actin filament nucleation via the pan-formin inhibitor SMIFH2 ^30^, cells expressing Rac1^P29S^ exhibited sustained proliferation under MAPKi treatment (Fig. 4i, j, Extended Data Fig. 2g), indicating that Rac1^P29S’^s proliferative advantage is driven specifically by the dendritic actin network formed by Arp 2/3 activity. Among regulators of cell proliferation, the protein product of the tumor suppressor gene NF2, Merlin, is associated with lamellipodia^31^, and thus may be a candidate for establishing a link between dendritic actin polymerization and enhanced cell cycle progression. Merlin is phosphorylated at Serine 518 by p21-activated kinase PAK1 ^31^, which also localizes to regions of dendritic actin polymerization ^32^. Phosphorylation at Serine 518 inactivates Merlin’s ability to suppress proliferation ^33^. Indeed, we observed that Merlin is highly phosphorylated in Rac1^P29S^ cells upon MAPKi treatment (Fig. 4k), consistent with the lamellipodia phenotype in MAPKi-treated Rac1^P29S^ cells and supporting a model where Rac1^P29S^ exhibits a proliferative advantage only under conditions that are otherwise not permissive for proliferation. Most critically, inhibition of Arp2/3 activity with CK666 abrogates Merlin phosphorylation in Rac1^P29S^ cells, while treatment with the control inhibitor CK689 does not (Fig. 4l). This underlines the requirement of Arp 2/3-dependent dendritic actin network assembly for Rac1^P29S’^s effect on sustaining cell proliferation under MAPK inhibition.

In summary, this work describes the role of the hyperactive Rac1^P29S^ mutation as a proliferative protection mechanism that confers drug resistance and metastatic advantage in melanoma (Extended Data Fig. 2h). The Rac1^P29S^ proliferative pathway is mechanosensitive and requires engagement of the cell with the ECM, but is independent of the canonical integrin-FAK-actomyosin axis co-noted with mechanically stimulated proliferation and survival ^20, 21^. The pathway bypasses MAPK signaling, explaining the clinically reported failure of BRAF targeting drugs in melanoma patients carrying the Rac1^P29S^ mutation ^34^. We also show in a preclinical model of melanoma progression that this pathway confers a proliferation advantage to metastatic cells in secondary sites, where they typically encounter a harsh, foreign microenvironment. Thus, we interpret the parallelism of growth suppression upon drug treatment and during metastasis as a convergence of these hallmark processes of aggressive cancers onto proliferation pathways sustained by Rac1. Unlike traditional oncogenes that are constitutively active, Rac1^P29S^ drives proliferation conditionally, conferring the greatest advantage to cells experiencing a growth challenge. Whether this adaptation is due to the disinhibition of a Rac1^P29S^ regulator or compensatory activation of the Rac1^P29S^ pathway remains to be examined. This along with the ERK-independence of this pathway opens the door to the development of alternative strategies for melanoma treatment, especially in view of the frequent alterations in Rac1-activating proteins in melanoma as well as other forms of cancer ^35^.

At the core of the pathway is the massively amplified, exponential growth of the dendritic actin network in cellular lamellipodia, which inactivates the tumor suppressor NF2/Merlin that has been shown to directly inhibit cell cycle progression through a number of pathways ^36–38^. Interestingly, linear growth of an actin network of straight filaments is insufficient to activate the pathway. While the ultrastructural details of the coupling between actin polymerization in a dense brush of filaments and Merlin inactivation remain to be uncovered, our data show a minimal requirement of the pathway for mechanical resistance by the plasma membrane and cell cortex against lamellipodial actin filament assembly: First, Rac1^P29S^-mediated drug resistance is completely abrogated in suspended cells, where no such resistance can arise. Second, for cells plated on stiffer substrates, where the plasma membrane/cortex system is less compliant to actin polymerization than on softer substrates, the proliferative advantage conferred by the pathway is enhanced. Consistent with this interpretation, under growth challenge by MAPK inhibitors, the engagement of the Rac1^P29S^ mediated pathway increases cell traction force, a direct measure of the force level generated by actin polymerization against the plasma membrane/cortex system ^39, 40^. Of note, the engagement of cells with the substrate does not require integrin-dependent matrix adhesions, but non-canonical and potentially even passive forms of cell-matrix coupling is sufficient to elicit the pathway. We point out that the mechanical induction of the pathway occurs on substrates with stiffness as low as 500 Pa. Thus, the mechanical engagement of cells with essentially any non-fluid tissue in the human body will suffice to instigate the pro-proliferative conditions prompted by Rac1-hyperactivation. This lends the suggestion that the same pathway can be active in micrometastatic tissues, where cell-cell adhesions may be equally abundant as cell-matrix adhesions. Hence, our study unveils a mechanism of cell growth that may be active in a wide range of scenarios, from development to neoplasm to homeostasis of challenged tissue.

## Methods

### Constructs and Reagents

Rac1 mutant and wild type viral vectors were generated from the pcDNA3-GFP-Rac1 wild type (Rac1^WT^) construct (Cell BioLabs). The QuikChange Lightning Mutagenesis Kit (210515, Agilent) was used to introduce the single-base pair 85C>T transition in the pcDNA3-GFP-Rac1^WT^ coding sequence resulting in the proline-to-serine mutation at amino acid 29 (Rac1^P29S^). Primers for the reaction were designed using the Agilent online tool (www.agilent.com/genomics/qcpd): CTGATCAGTTACACAACCAATGCATTTTCTGGAGAATATATCCC (forward) and GGGATATATTCTCCAGAAAATGCATTGGTTGTGTAACTGATCAG (reverse). Coding sequences of Rac1^P29S^ and Rac1^WT^ were cut out of the resulting pcDNA3-GFP-Rac1^P29S^ construct and the original pcDNA3-GFP-Rac1^WT^ construct, respectively using the enzymes EcoR1 and Xho1. These coding sequences were ligated into a pLVX-IRES-puromycin lentiviral construct (pLVX-puro) that was cut with the same enzymes to create pLVX-puro-Rac1^P29S^ and pLVX-puro-Rac1^WT^ constructs. Undigested pLVX-puro construct was included in subsequent steps to serve as an appropriate empty-vector negative control to rule out effects due to expression of the backbone vector. HEK293T cells were transfected using PEI at 3ul/ug total DNA concentration with these constructs (5ug) and VSV envelope vector pspax2 (7ug) and pmd2g viral packaging construct (5 ug). Viral media was harvested, filtered (0.45 um), and mixed with 1ul/ml dilution of polybrene. To generate A375 cells lines that exogenously express Rac1^WT^ (+WT), Rac1^P29S^ (+P29S), and empty pLVX-puro vector (+EV), parent A375 cells were spin-infected for 1hr at 3000 rpm. Viral media was replaced with fresh media 24 hrs following infection. Cells transduced with pLVX-puro-Rac1^WT^, pLVX-puro-Rac1^P29S^, and undigested pLVX-puro (empty vector) were selected with 2ug/ml puromycin.

To titrate expression of fluorescent protein-tagged constructs and avoid overexpression artifacts, a series of truncated CMV promoters were created using the pLVX-shRNA2 backbone (Clontech). Briefly, the U6 promoter, multiple cloning site, and ZsGreen reporter gene were removed and replaced with iteratively increasing 100 basepair increments of the CMV promoter starting with the 5’ end. Thus, 6 constructs were prepared, with the first 100, 200, 300, 400, 500, and 600 bp of the promoter region, fluorescent protein fusions were cloned downstream using the restriction enzymes SpeI and XhoI. These constructs will be made available on Addgene. To label adhesions, mNeonGreen-Paxillin-22 (Allele Biotechnology), a C-terminal fluorescent protein fusion with a 22 amino acid linker, was cloned into the 100 bp variant. To label the actin cytoskeleton, mNeonGreen-Actin-C-35 (Allele Biotechnology), an N-terminal fluorescent protein fusion with a 35 amino acid linker, was cloned into the 300 bp variant. Lentiviral particles for both constructs were prepared according to the manufacturers recommendations, and infected A375 cells were enriched with fluorescence activated cell sorting (FACS) using a FACSAria system (Children’s Research Institute Flow Cytometry Facility).

### Xenograft tumor model and Immunohistochemistry

Viral supernatant prepared from a dsRed2-P2A-Luc lentiviral construct was provided generously by the Morrison lab to generate dsRed-luciferase positive cells for xenograft tumor injections and subsequent bioluminescence imaging to detect metastases ^41^. Following infection, dsRed positive cells were enriched using flow cytometry and expanded. 100 cells were counted and prepared in 25% Matrigel as described in ^41^ for subcutaneous injection into the flank region of NSG mice. Tumor growth was measured using a Marathon CO030150 Digital Caliper until tumors reached around 2.0 cm but no more than 2.5 cm before mice were euthanized for end-point analysis of metastasis. Bioluminescence imaging of dissected organs and analysis were performed as described in ^41^. These experiments were performed according to the protocol approved by the UT Southwestern Institutional Animal Care and Use Committee (protocol 2016-101360). Dissected lungs were fixed in 10% neutral-buffered formalin for 48 hrs on a rotator at room temperature then transferred to PBS. Tissues were paraffin embedded and sectioned by the UT Southwestern Histo Pathology Core. Immunohistochemistry was performed to dual label mouse lung tissues for melanoma metastases using rabbit anti-RFP antibody (Rockland) to recognize dsRed-luciferase positive disseminated human melanoma cells and mouse anti-S10-phospho-Histone H3 (9706, Cell Signaling) antibody to recognize mitotic cells. Briefly, sections were deparaffinized for 5 min in a xylene bath, done thrice. Sections were hydrated in sequential ethanol baths: 100% ethanol, 5 min, 90% ethanol, 2 min, 80% ethanol, 2 min, 70% ethanol, 2 min, and 50% ethanol, 2 min. Sections were left under a gentle stream of running water for 5 min. Antigen presentation was achieved by cooking sections in sodium citrate buffer (10 mM sodium citrate, 0.05% Tween 20, pH 6.0) inside a heated pressure cooker for 5 min. Following cool down, slides were rinsed with water and removed to 1x TBST. To block endogenous peroxidase, slides were incubated with a 3% H2O2 solution for 15 min. Sections were rinsed for 5 min twice with 1x TBST and Biotin-Avidin blocking was performed (SP-2001, Vector Laboratories). The Vector M.O.M Peroxidase-based Immunodetection Kit (Vector Laboratories) was followed for additional blocking, mouse anti-S10-phospho-Histone H3 primary antibody incubation (1:300 dilution, 4°C, overnight), and secondary antibody incubation. Phospho-Histone H3 positive tissue was visualized using the ImmPACT DAB Peroxidase Substrate Kit (SK-4105, Vector Laboratories). Sections were washed with 1x TBST then blocked and incubated in rabbit anti-RFP antibody (Rockland) according to the ImmPRESS-AP Reagent Anti-Rabbit IgG alkaline phosphatase-based staining kit (MP-5401, Vector Laboratories). Metastatic melanoma cells were visualized using ImmPACT Vector Red alkaline phosphatase substrate (SK-5105, Vector Laboratories). Sections were then hematoxylin stained (15 sec, H3404, Vector Laboratories). To quantify proliferation in metastases, Ilastik pixel classification was used to determine pixel area of all metastatic nodules positively labeled for dsRed expression. All metastatic nodules with pixel areas between 200,000 and 400,000 were randomized and phospho-Histone 3 positive nuclei were hand counted to determine number of proliferating cells (n ≥ 9 metastatic lung nodules for each condition).

### Cell Culture and Proliferation Assays

A375 (CRL-1619, ATCC), WM3060 (WC00126, Coriell Institute), 451Lu (WC00059, Coriell Institute), and IGR1 (ACC236, DSMZ) cells were cultured using DMEM (Gibco) supplemented with 10% FBS and 0.2% antibiotic-antimycotic (15240062, Gibco). Fraction of proliferating cells was determined using the Click-iT EdU Alexa Fluor Imaging Kit (Molecular Probes). Briefly, 15,000-30,000 A375 cells or 70,000 IGR1 cells were seeded into each well of a 12-well plate and drug treated for 48hrs. Cells were incubated with 10uM of EdU for 3hrs prior to fixation with 4% PFA and permeabilization with 0.5% Triton. Edu-positive nuclei were labeled according to manufacturer protocol followed by labeling of all nuclei with Hoechst (10mg/ml) diluted 1:2000. An inverted phase contrast and fluorescence Nikon Ti-Eclipse microscope equipped with a Zyla sCMOS camera, SOLA solid state white-light excitation system and a motorized filter turret with filters for DAPI, FITC and TRITC along with Nikon Elements image acquisition software was used to image Hoechst and Edu-positive nuclei at 10x magnification. For all Edu incorporation assays, three biological replicates were performed on separate days. Around five images, sampling multiple regions of the well, were acquired for each condition (n≈15).

Nuclei were counted automatically in a custom pipeline for image processing, quality control and condition analysis developed in Matlab (The MathWorks, Inc.). The detection algorithm uses the 99 percentile of the background noise to estimate an adaptive threshold for significant nuclei signal. In order to discriminate nuclei in close proximity in the mask of significance, local intensity maxima are detected on the band-passed filtered image before watershed segmentation. The band-pass filter allows for selectivity in scale of the object of interest. The scales used for the band-pass filter are 8 pixels and 15 pixels for low-pass and high-pass filter respectively. Since we didn’t observe variations in nuclei size across experiment, this algorithm provides us with an unbiased method to compare nuclei accounts across conditions. We then used the u-track GUI ^42^ to allow for a systematic review of the stationarity of imaging condition. The measurements associated to each nuclei detections (position, intensity, channel and time) are then grouped by condition and stored in a spreadsheet for flexible plotting.

For experiments testing proliferation sensitivity to drug for cells cultured in low adhesion or soft collagen, percent proliferation was determined using the Click-iT Plus Edu Alexa Fluor Flow Cytometry Kit (C10632, Molecular Probes). For low adhesion experiments, 300,000 cells were seeded into tissue culture coated 6 well plates or ultra-low attachment 6 well plates (Corning) to keep cells in suspension. For soft collagen experiments, 300,000 cells were counted and resuspended in 2 mg/mL collagen (Advanced Biomatrix) (1 mL cocktail of pre-warmed reagents: 100 uL 10x PBS, 10uL 1M NaOH, 250 uL H2O, 640 uL 3mg/mL collagen). 100 mM ribose (R9629, Sigma) was supplemented in the collagen cocktail to increase rigidity by enhancing crosslinking ^24^. Cells were plated in dishes prewarmed to 37°C. Following drug treatment and Edu incubation, cells were either trypsinized from plastic dishes, collected from suspension dishes by aspiration, or released from collagen gels by incubation in collagenase (1mg/ml in PBS, added 1:1 collagen to collagenase volume) for 3 hrs at 37oC (5030, Advanced Biomatrix) prior to neutralization with media then fixation and permeabilization according to manufacturer’s protocol. Labeled cells were analyzed using a FACSAria II Cell Sorter (BD Biosciences, San Jose, CA) and FlowJo v10 analysis software.

### Western blotting and Activity Assays

Cells were washed twice in chilled Hank’s Balanced Salt Solution containing calcium and magnesium (HBSS, 14025092, Gibco) and lysed in chilled 1x JS buffer (2x JS: 100mM Hepes, pH 7.5, 300mM NaCl, 10mM EGTA, 3mM MgCl2, 2% Glycerol, 2% Triton X-100, supplemented with protease and phosphatase inhibitors). Protein concentrations were determined using Pierce BCA Assay kit (Thermo). Samples were run on 4-20% Mini-PROTEAN pre-cast gels (Biorad) or either 10% or 15% homemade gels and transferred to PVDF membranes (Thermo). For the Rho activity assay GST-RBD (Rhotekin) was purified as previously described (Isogai et al., 2015 JCS). The RBD assay was carried out by incubating 500 μg of total cell lysate (lysis buffer: 50mM Tris pH 7.5, 150 mM NaCl, 10mM MgCl2, 1% Triton X-100, 1mM DTT, supplemented with protease and phosphatase inhibitors) with 30 μg of GST-RBD and GST-CRIB, respectively, as previously described (Isogai et al., 2015 JCS). Antibodies used and dilutions are as follows: T202/Y204-pERK 1:500 (E10, Cell Signaling), ERK 1:1000 (sc-93, Santa Cruz), pMEK 1/2 1:1000 (9121, Cell Signaling), MEK 1/2 1:1000 (9122, Cell Signaling), Cyclin D1 1:500 (sc-8396, Santa Cruz), GAPDH 1:5000 (G9545, Sigma), Y397-pFAK 1:500 (D20B1, Cell Signaling), FAK 1:1000 (D2R2E, Cell Signaling), Y118-pPaxillin 1:1000 (Cell Signaling), Y31-pPaxillin 1:1000 (R&D Systems), Paxillin 1:10,000 (ab32084, Abcam), Talin 1:1000 (Sigma), β-Actin 1:100,000 (AC15 Sigma), RhoA 1:500 (Cell Signaling), T18/S19-pMYLC2 1:1000 (Cell Signaling), MYLC2 1:1000 (D18E2, Cell Signaling), JNK 1:500 (sc7345, Santa Cruz), pJNK 1:1000 (sc6254, Santa Cruz), S518-pNF2 1:100 (9163, Cell Signaling), NF2 1:1000 (D3S3W, Cell Signaling). Western blot quantification was done using the gel densitometry application in ImageJ.

### Immunofluorescence

Paxillin in cell adhesions was visualized using immunofluorescence at least 24 hrs following cell plating on #1.5 acid-etched coverglasses in 6-well plates or #1.5 glass-bottom chamber slides (155382, nunc). Cells were washed thrice in PBS then incubated for 2 min in a 2% paraformaldehyde/0.5% Triton X-100 solution followed by 4% paraformaldehyde for 30 min. Samples were washed thrice in PBS. Quenching-PBS (Q-PBS, 225 ml PBS, 22.5 mg saponin, 4.5 g BSA, 225 mg lysine, pH 7.4) was added to samples for 30 min to block non-specific binding sites. Rabbit anti-paxillin antibody 1:500 (ab32084, Abcam) was prepared in Q-PBS and added to samples for 1 hr then washed for 5 min, 6 times in Q-PBS. Samples were incubated in anti-rabbit Alexa 488-conjugated secondary antibody 1:500 (A11034, Molecular Probes) prepared in QPBS for 1hr, and washed for 5 min, 6 times in PBS. Samples were post-fixed for 10 min in 4% paraformaldehyde and washed thrice in PBS. Paxillin-labeled adhesions were visualized with total internal reflection fluorescence microscopy (TIR-FM) using a Nikon Ti-Eclipse inverted microscope equipped with a 100x 1.49 NA Apo TIRF objective, a Diskovery Platform (Andor Technology) for TIR-FM illumination, and an sCMOS camera.

### Live cell imaging

Cell were plated in 35 mm WillCo dishes on #1.5 cover glasses coated with thin (~30 µm) layers of soft silicone gels with high refractive indices, making the substrates both deformable under cell traction forces and compatible with TIRF microscopy ^43^. Surfaces of the gels were decorated with covalently bonded 40 nm dark red (660/680 nm) fluorescent beads (ThermoFisher), which served as tracer particles, enabling traction force microscopy (TFM). Silicone gel substrates were coated with fibronectin by incubating them for 30 minutes under a solution of 20 ul of a 10 mg/mL 1-ethyl-3-(3-dimethylaminopropyl) carbodiimide hydrochloride (EDC) solution, 30 ul of a 5 mg/mL fibronectin solution, and 2 mL of Ca^2+^ and Mg^2+^ containing Dulbeccos Phosphate Buffered Saline (DPBS, 14040117, Gibco). Following coating, substrates were rinsed multiple times with DPBS followed by a final rinse in DMEM and incubated at 37°C prior to cell plating and drug treatment (0.003% DMSO, 33.3nM Dabrafenib). High-resolution imaging was performed 48 hrs later using a DeltaVision OMX. For traction-force microscopy, the microscope was operated in a ring-TIRF illumination condition, which permits more homogeneous illumination of the basal surface of the cell. Images were acquired at a 60x magnification, providing an 80 nm pixel size and Nyquist sampling. Paxillin and the fluorescent tracer particles were imaged with 488 and 640 nm lasers, respectively, and focus offsets were introduced for each spectral channel to maximize image focus. Following imaging, cells were removed rapidly by adding 1 mL of 30% bleach to the 2 mL of cell media, and the tracer particles on the substrate at each cell position were imaged under the relaxed condition. For actin imaging, cells were plated in #1.5 glass-bottom chamber slides (155382, nunc), and the microscope was operated in an oblique illumination mode, thus minimizing unnecessarily illumination of the cells. Images were acquired every 5 seconds, which is sufficient to Nyquist sample actin polymerization, protrusion, and retraction events, in time.

### Image Analysis

Traction force was analyzed using a Matlab-based algorithm that uses L1 regularization ^25^. The regularization parameter was determined by optimal regularization selection criteria - an inflection point in the L-curve, which is a plot between the residual norm vs. the self-norm of the traction solution. The strain energy (1/2 * displacement * traction), which represents the overall mechanical work done by a cell on the soft gel, was quantified from traction maps and displacement maps of individual cells, integrated over an entire, segmented cell area. Strain energy distributions of A375 cells expressing empty vector, Rac1^WT^, or Rac1^P29S^ were compared following treatment for 48hrs with either 0.003% DMSO or 33.3nM Dabrafenib.

Focal adhesion (FA) segmentation was performed as described previously ^25^, based on a combination of Otsu and Rosin thresholding after preprocessing images with noise removal and background subtraction. Based on the segmentation, statistics for FA area were determined and compared for A375 cells expressing empty vector, Rac1^WT^, or Rac1^P29S^ following treatment with either 0.003% DMSO or 33.3nM Dabrafenib

### CRISPR/Cas9 genome editing

For the single-base pair change of endogenous Rac1^P29S^ in IGR1 cells to Rac1^WT^, CRISPR/Cas9 mediated genome-editing was used, and homologous recombination was achieved using a ssODN repair template by following the protocol described by the Zhang Lab ^44^. The sgRNA guide sequences 5’-TACACAACCAATGCATTTTC-3’ and 5’-ATATTCTCCAGAAAATGCAT-3’ were designed using the CRISPR Design Tool (http://tools.genome-engineering.org) and the ssODN repair sequences AAGATACTTACACAGTAGGGATATATTCTCCAGGAAATGCATTGGTTGTGTAACTGATCAGTAGGCAAGT and AAAACTTGCCTACTGATCAGTTACACAACCAATGCATTTCCTGGAGAATATATCCCTACTGTGTA AGTAT corresponding to each guide, respectively, were designed using the construct visualization tool, Benchling.com. Guide sequences were cloned into the pSpCas9(BB)-2A-GFP (PX458) vector (48138, Addgene) ^44^. IGR1 cells plated in 10cm dishes were transfected with both guides independently following 30 min treatment of cells with 0.05 uM of SCR7 pyrazine (SML1546, Sigma) using 15 ug of CRISPR/Cas9-guide construct DNA, 30 ul of ssODN reconstituted to 10 uM and 30 ul of Lipofectamine LTX Reagent (15338030, Thermo). GFP-positive cells were then plated as single cells using FACS into 96-well plates containing IGR1-pre-conditioned media. Colonies were expanded in 24 well plates and screened for successful genome-editing using gDNA extraction using QuickExtract (QE09050, Epicentre) and Sangar sequencing.

For CRISPR/Cas9-mediated knockout of ARPC1B, selected DNA target sequences from exon
3 of ARPC1B were pasted into the CRISPR design tool CRISPOR (http://crispor.tefor.net). Resulting potential target sites with a high efficiency score were used for designing the sgRNA constructs (20 nucleotides), which were cloned into pSpCas9(BB)-2A-GFP (pX458; Addgene plasmid ID: 48138) using BbsI and sequence verified ^44^. To disrupt ARPC1B gene expression, a blasticidin selection cassette was inserted at the cut site employing a selfcleaving donor vector. The sequence for the sgRNA used to target ARPC1B is as follows: 5’-CTCGTGCACCTTGGTCCATT-3’. ARPC1B knockout and the Arp 2/3 complex were evaluated with western blots using the following antibodies: anti-ARP3 1:500 (sc-48344), anti-ARPC2 1:500 (sc-515754), and anti-ARPC1B 1:500 (sc-137125) all from Santa Cruz and anti-ARPC1A 1:1000 (HPA004334, Sigma).

### Data and Code Availability

The images and data generated during this study are available from the corresponding author upon reasonable request.

Custom codes used for data analysis include the current version of u-track particle tracking software (Version 2.2.1) and traction force microscopy software (Version 1.1.3), which are both are freely available for download from the Danuser Lab website http://www.utsouthwestern.edu/labs/danuser/software/).

## Acknowledgements

We would like to thank Dr. Sean Morrison for providing guidance for *in vivo* experiments and for critically reading the manuscript. We are grateful for assistance from the UT Southwestern Histo Pathology Core for providing guidance and services for tissue handling and also generous access to supplies and equipment. We are also grateful to the Shay/Wright Lab for both resources and guidance: Dr. Andrew Ludlow, for input on CRISPR/Cas9 genome editing, Crystal Cornelius for managing and sharing the pGIPZ shRNA construct library, and Krishna Luteal for his immunostaining protocol. We would also like to thank Michael Abrams and the Alto Lab for guidance for CRISPR/Cas9 genome editing and Dr. Marcel Mettlen in the Schmid Lab for his immunofluorescence protocol. This work was supported by funding from the following grants: NIH F30 CA206399 (A.M.), NIH F32 GM117793 (K.M.D.), Human Frontier Science Program (P.R.), NIH K25 K25CA204526 (E.S.W.), CPRIT R1225 (G.D.) and NIH R01 GM071868 (G.D.).

## Author Contributions

A.M. and G.D. conceived the project. E.W. provided intellectual input. A.M. E.W. and G.D. wrote the manuscript and A.M made the figures. A.M. designed and performed the experiments and analysis. K.D., T.I., and V.M. performed experiments. K.D. provided all constructs for imaging. S.K. provided intellectual and technical input for mouse studies. T.I. provided the ARPC1b knock out cell line. S.H. performed adhesion and TFM analyses and P.R. implemented the automatic pipeline for counting proliferating cells. Silicone substrates were provided by A.G. All authors reviewed and provided feedback on the manuscript.

## Competing Interests

The authors declare no competing interests.

---

**Extended Data Figure 1.**
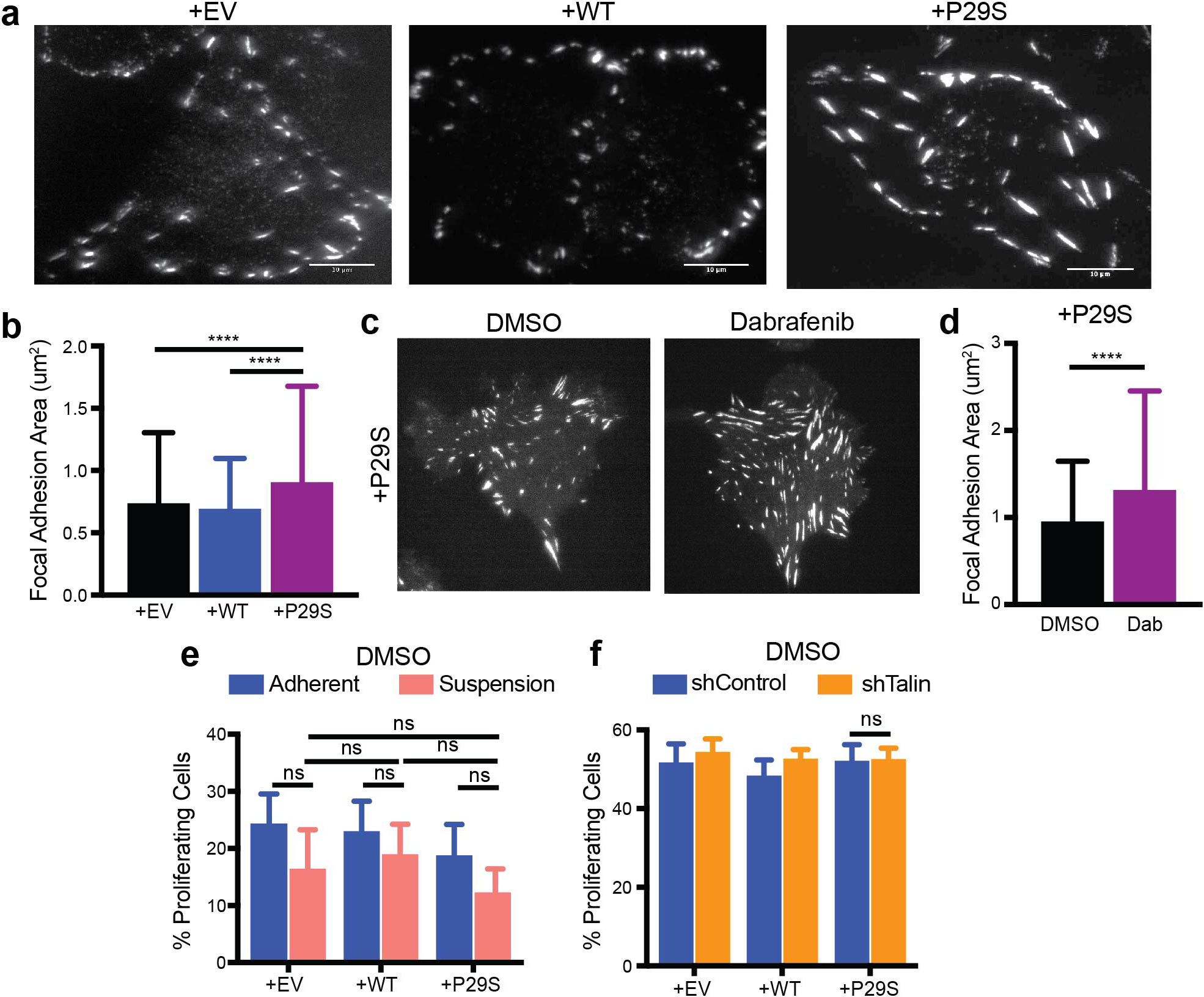
**a,b** Differences in focal adhesions between A375 cells expressing Rac1P29S (+P29S), Rac1^WT^ (+WT), or empty vector (+EV). (**a**) Immunofluorescence of Paxillin. (**b**) Quantification of adhesion size. **c,d,** Focal adhesions in +P29S A375 cells upon 48 hr DMSO (0.003%) and Dabrafenib (33.3 nM) treatment (**c**) visualized by live cell fluorescence imaging of mNeonGreen-tagged Paxillin and (**d**) quantification of adhesion size. **e,f** Proliferation assay applied to A375 cells expressing +EV, +WT, and +P29S cultured under DMSO treatment (0.003%) (**e**) on plastic versus low-adhesion dishes and (**f**) under stable expression of shControl and shTalin constructs. To compare adhesion areas, a Tukey’ s multiple comparison’ s test was used in (**b**), while an unpaired *t*-test was used in (**d**). A 2-way ANOVA and a Tukey’ s multiple comparison’ s test was used to determine p-values in (**e**) to determine differences between cell lines grown in suspension: p-values, +EV vs. +WT: 0.7980, +EV vs. +P29S: 0.5506, and +WT vs. +P29S: 0.2312 and unpaired *t*-tests were used to compare for each cell line proliferation differences due to adherent vs. suspension culture: p-values, +EV: 0.1240, +WT: 0.3347, +P29S: 0.1145. A two-sample equal variance *t*-test was performed to determine significance in (**f**) for +P29S cells, p-value, shControl vs. shTalin: 0.741. All bar graphs represent mean ± s.d. ****P<0.0001.

**Extended Data Figure 2.**
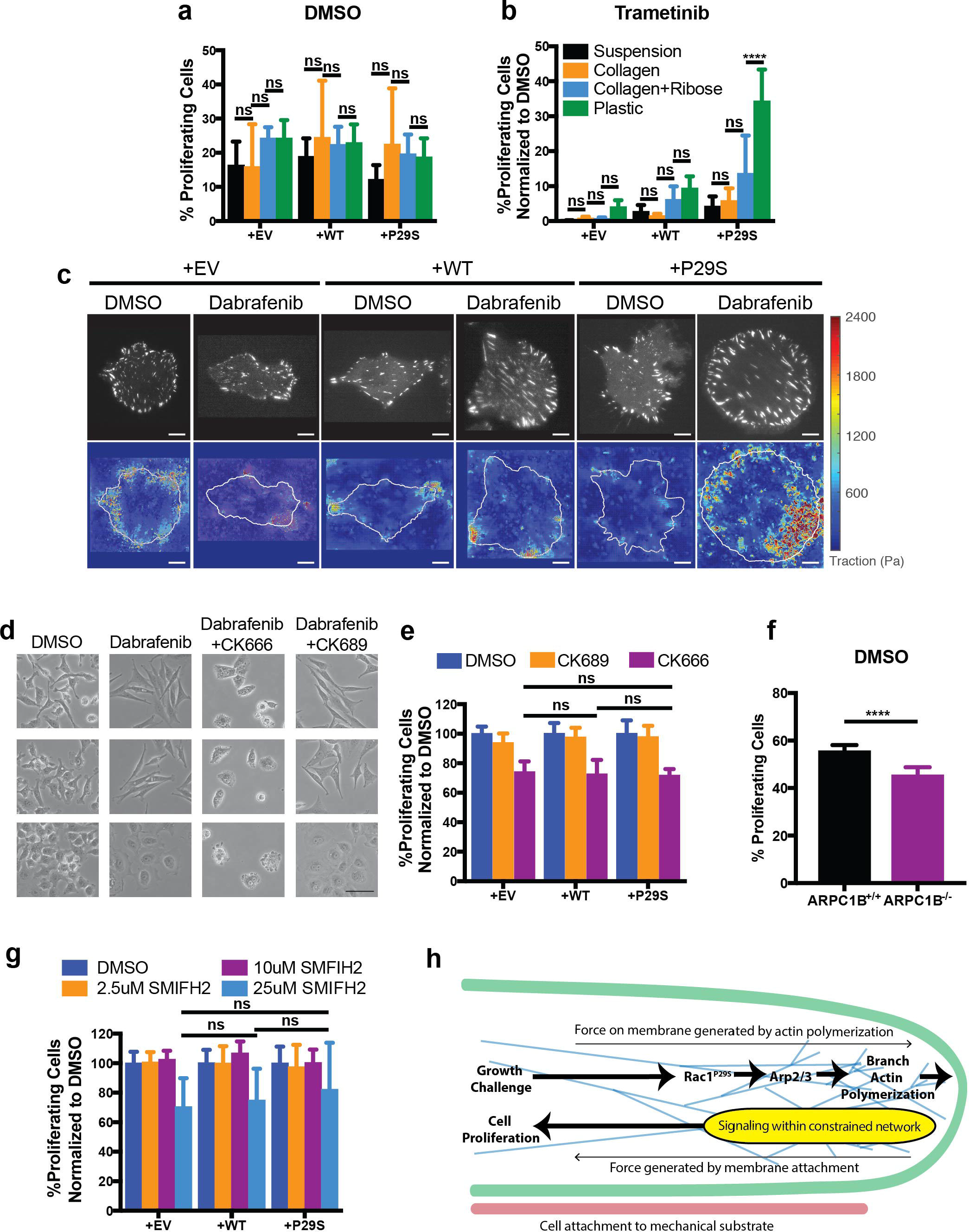
**a,b** Proliferation assay applied to respective A375 cells treated with DMSO (0.003%) or Trametinib (3.3 nM) and cultured using substrates of different stiffness. **c,** Traction force maps for respective A375 cells treated with DMSO (0.003%) or Dabrafenib (33.3 nM). Scale bar = 10 um. **d,** Cell morphology changes upon DMSO (0.003%) and Dabrafenib (33.3 nM) treatment and Dabrafenib treatment (33.3 nM) combined with CK666 (200 uM) or CK689 (200uM) as indicated. Scale bar = 50 um. **e-g,** Proliferation of cells upon DMSO (0.003%), CK666 (200 uM), CK689 (200uM), and SMIFH2 treatment. Treatment of cells with CK666 (**e**) and SMIFH2 (25 uM) (**g**) suppresses proliferation consistently regardless of Rac1 status. ARPC1B knockout (**f**) results in a small but significant drop in baseline proliferation in +P29S cells. Statistical significance was determined using a two-way ANOVA and Tukey’ s multiple comparisons test for (**a, b, e, and g**). An unpaired *t*-test was used to determine significance in (**f**). Cells listed as +P29S, +WT, and +EV are A375 melanoma cells exogenously expressing Rac1^P29S^, Rac1^WT^, and empty vector, respectively. **h,** Schematic summary showing Rac1^P29S^ driven cell cycle progression via mechanosensitive dendritic actin polymerization. All bar graphs represent mean ± s.d. ****P<0.0001.

**Supplementary Figure 1.**
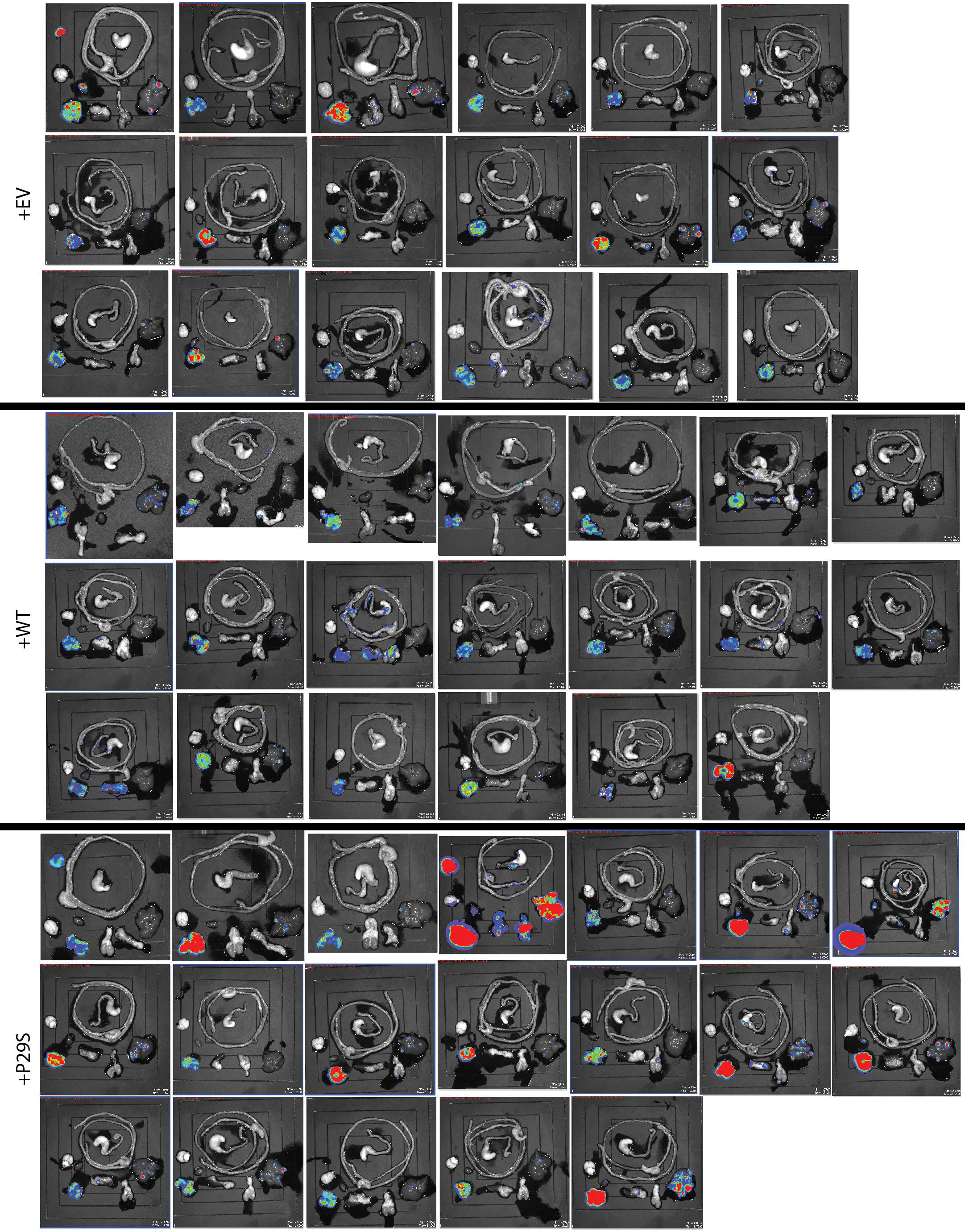
BLI signal detected in organs harvested from mice with primary tumors between 2.0-2.5 cm was used to measure metastatic burden.

**Supplementary Figure 2.**
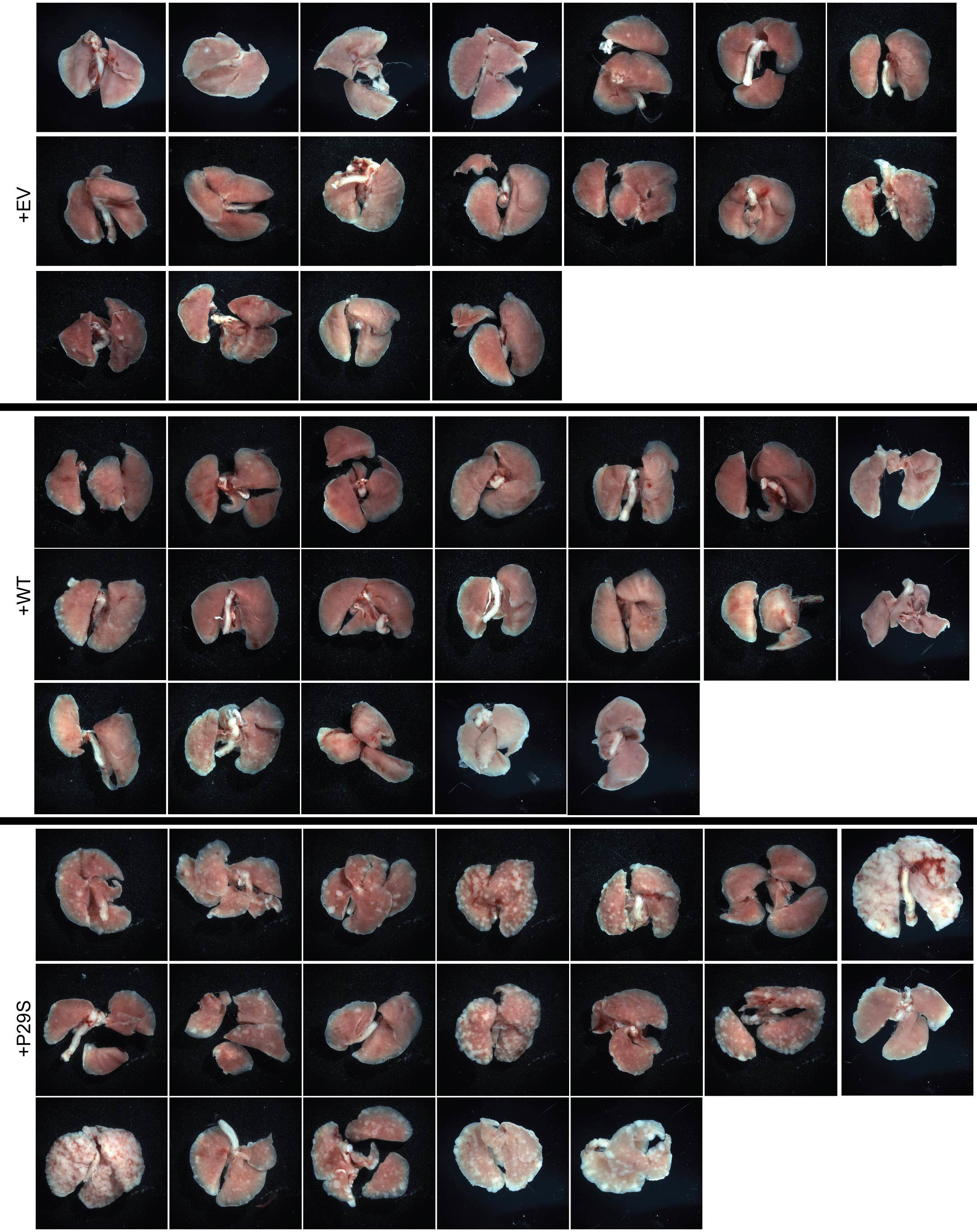
Macrometastases in gross lungs post fixation in 10% neutral-buffered formalin for 48hrs.

**Supplementary Figure 3.**
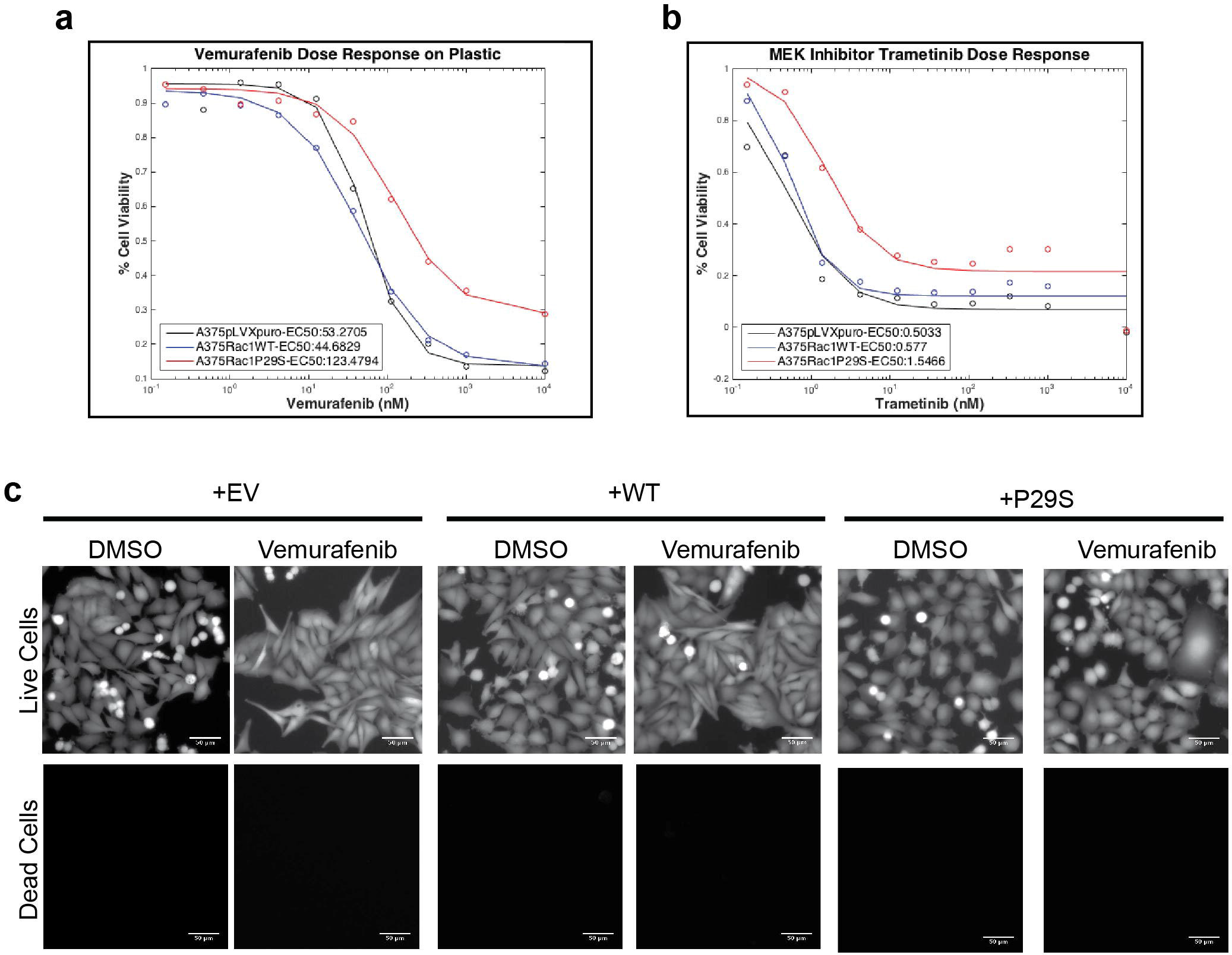
**a,b,** Our data reproducing the tetrazolium salt-based cell viability assay (CCK-8, Dojindo) used in Watson, et al. Cancer Research. 2014. that suggests that expression of Rac1^P29S^ can confer a survival and/or growth advantage upon MAPKi treatment. **c,** Calcein and ethidium homodimer labeling of live and dead cells, respectively, upon 48hr MAPKi treatment with 333.3 nM Vemurafenib reveals minimum cell death for all three cell lines (+EV, +WT, and +P29S), suggesting that the advantage conferred by +P29S expression observed in (**a**) is unlikely to be a survival advantage protecting against drug-induced cell death. Scale bar = 50 um.

**Supplementary Figure 4.**
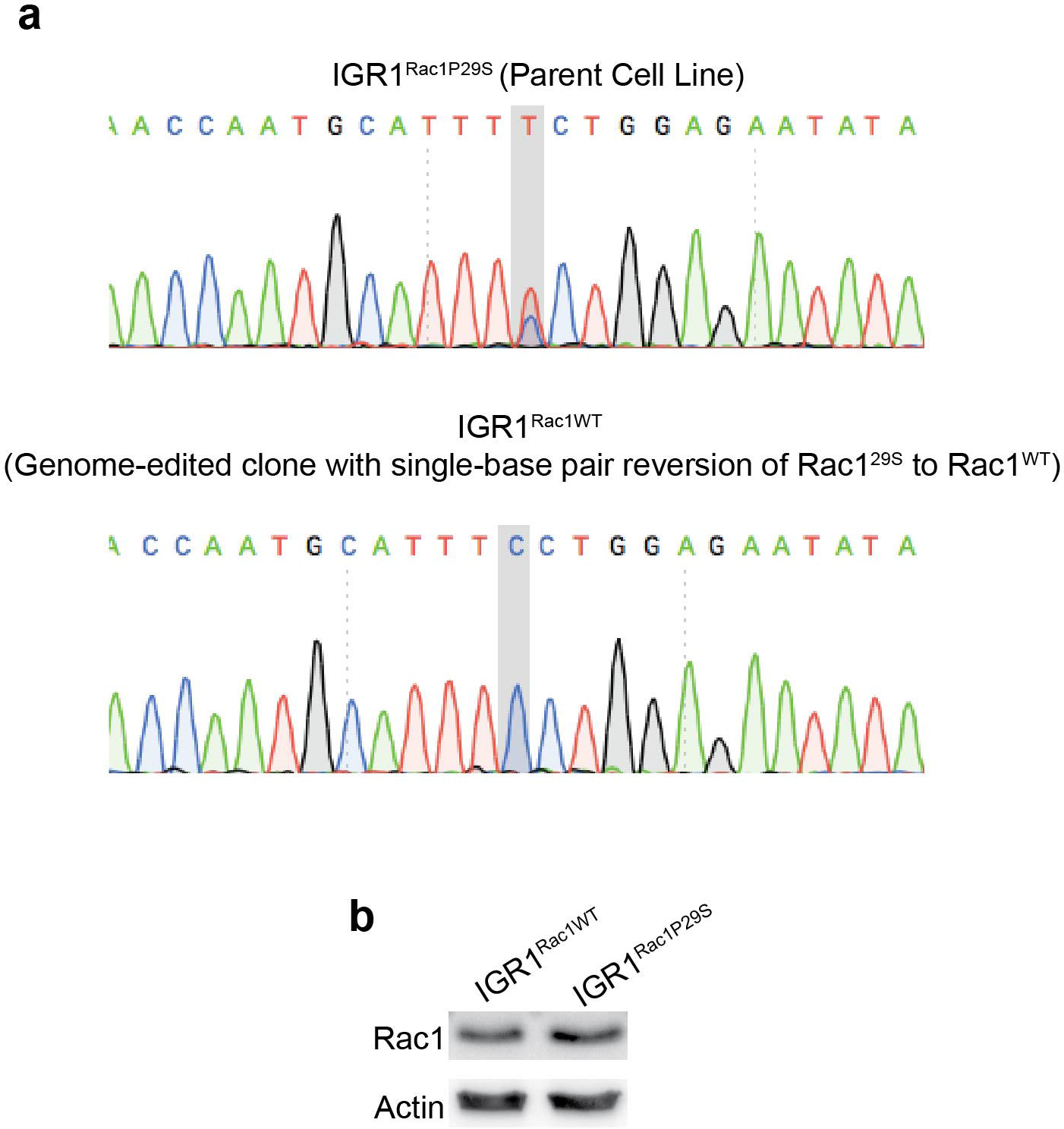
**a,** Sequencing chromatograms showing Rac1^P29S^ mutation present in parent IGR1 cells and the CRISPR/Cas9 mediated reversion by homologous recombination of the single 85C>T base pair transition back to Rac1^WT^. **b,** Immunoblotting confirms that parent and genome edited IGR1 cells express comparable levels of Rac1.

**Supplementary Figure 5.**
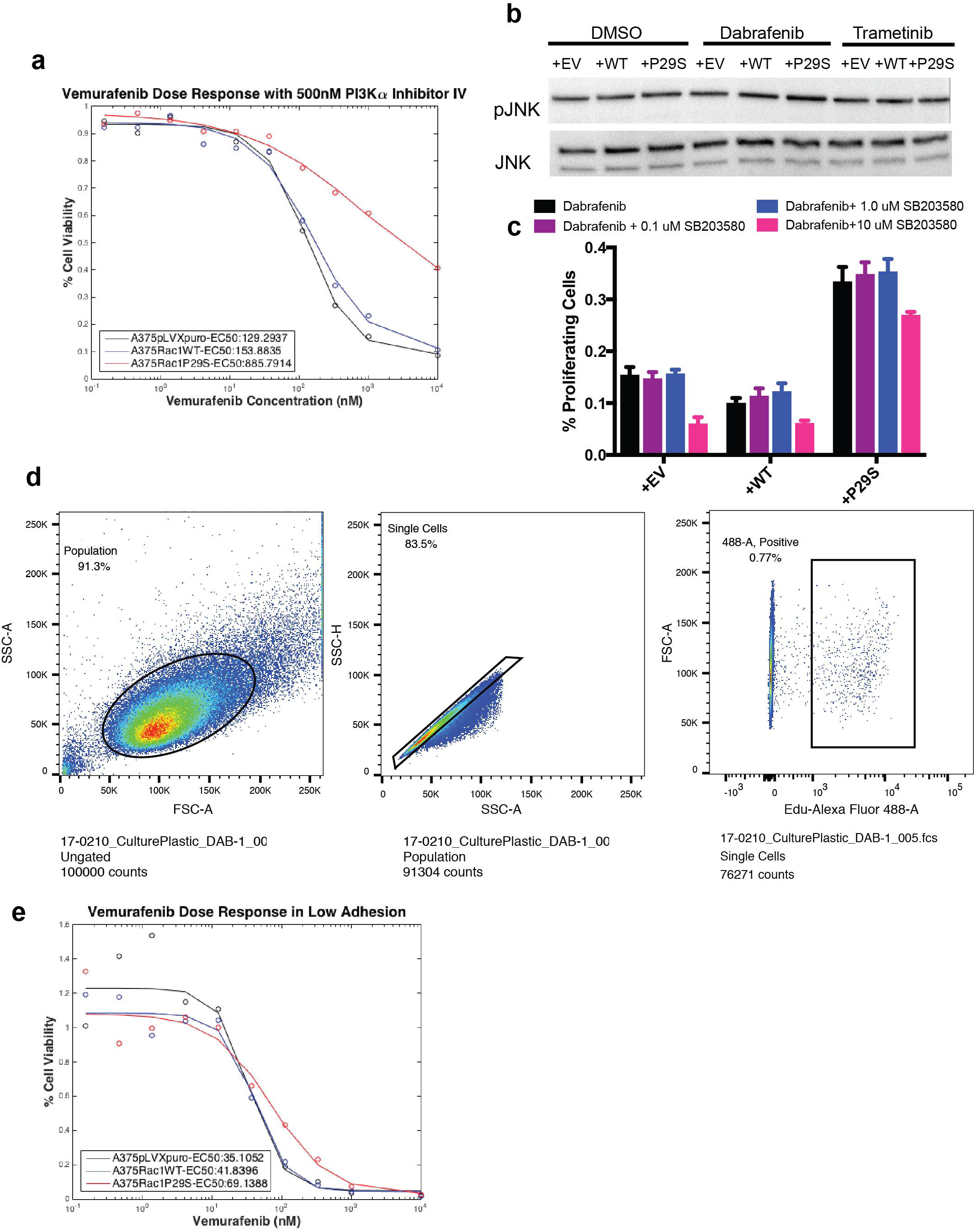
**a, b, c** Three alternative pathways tested for involvement in sustaining proliferation in +P29S cells upon MAPKi treatment. (**a**) Tetrazolium salt-based cell viability assay used to generate dose response curves for Vemurafenib at concentrations 0.15 nM, 0.45 nM, 1.37 nM, 4.11 nM, 12.3 nM, 37.0 nM, 111.1 nM, 333.3 nM, 1000 nM, 10,000 nM combined with inhibition of the PI3K pathway with 500nM PI3K alpha inhibitor IV. +P29S cells maintain growth advantage compared to +EV and +WT cells with PI3K inhibition. (**b**) Immunoblotting of JNK activity in +EV, +WT, and +P29S cells shows that JNK is not differentially regulated in +P29S cells. (**c**) Proliferation assay used to test effect of p38 MAPK inhibition with drug SB203580 on +P29S sensitivity to Dabrafenib. Control cells’ and +P29S cells’ Dabrafenib sensitivity appear to be comparably affected at all concentrations, suggesting p38 MAPK does not account for the +P29S-specific proliferative advantage observed. **d,** Flow cytometry plot for a single experimental condition (+EV cells, 33.3 nM Dabrafenib-treated, adherent culture) exemplifying gating strategy used to determine cell proliferation for assays using Click-iT Plus Edu Alexa Fluor Flow Cytometry Kit (C10632, Molecular Probes). **e,** Vemurafenib dose response curve from tetrazolium salt-based assay as described in (**a**) for cells cultured in low-adhesion dishes. +P29S cells have sensitivity to MAPKi that appears comparable to control cells upon growth in suspension.

**Supplementary Figure 6.**
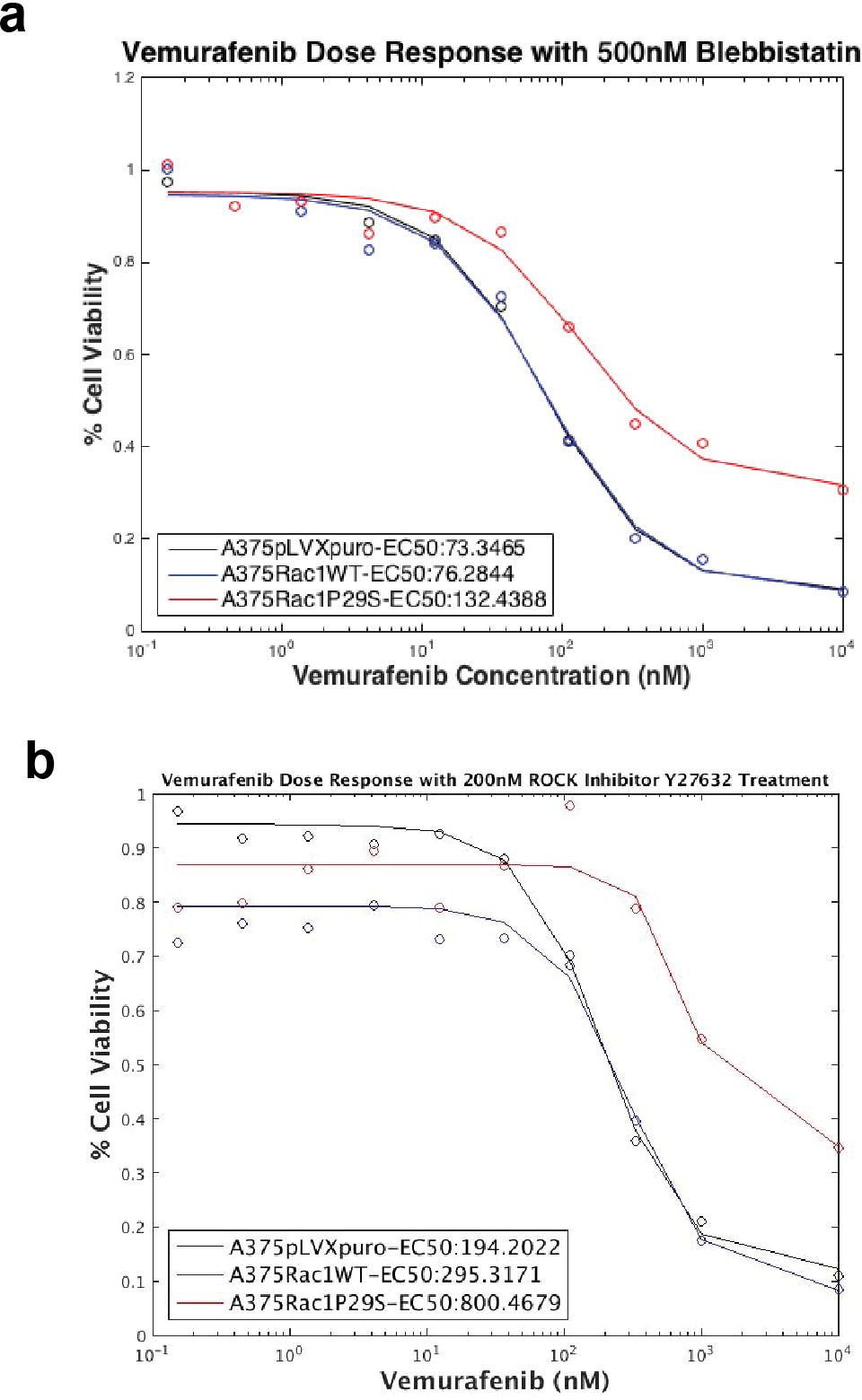
**a, b,** Tetrazolium salt-based cell viability assay used to generate dose response curves for Vemurafenib at concentrations 0.15 nM, 0.45 nM, 1.37 nM, 4.11 nM, 12.3 nM, 37.0 nM, 111.1 nM, 333.3 nM, 1000 nM, 10,000 nM combined with inhibition of (**a**) myosin with 500nM Blebbistatin and (**b**) Rho kinase (ROCK) with 200nM Y27632. +P29S cells retain growth advantage during MAPKi treatment when cell actomyosin contractility is inhibited.

**Supplementary Figure 7.**
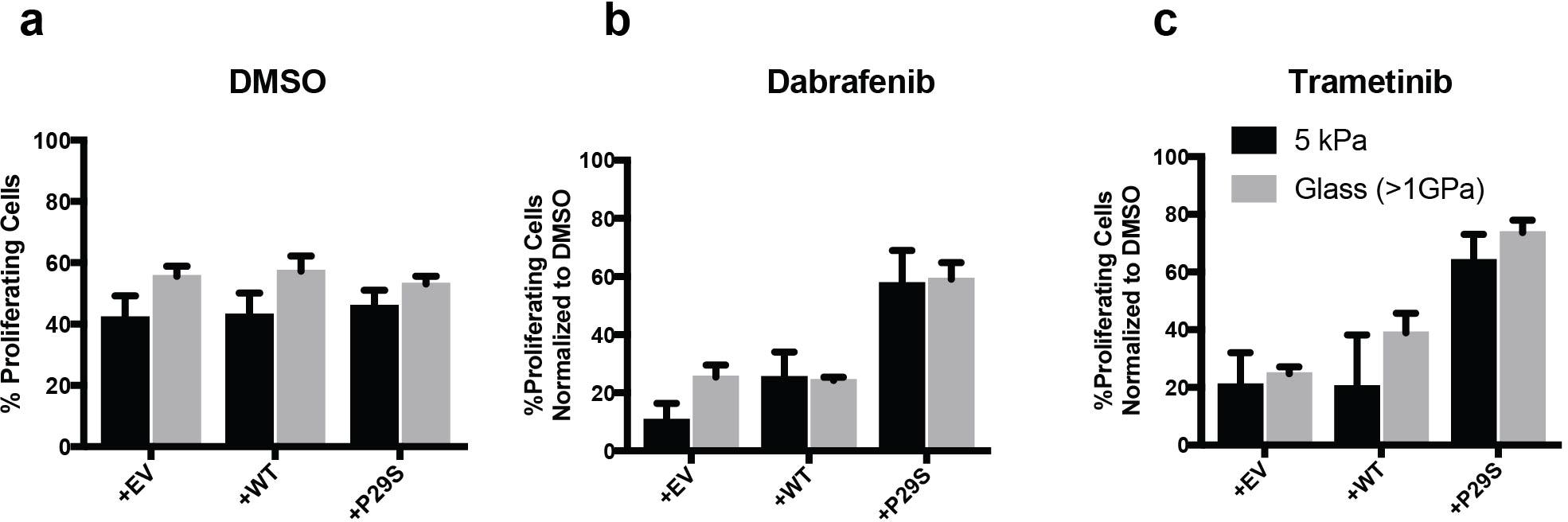
**a, b, c,** Proliferation assay showing effect of substrate stiffness 5 kPa versus glass (>1GPa) on sensitivity of +EV, +WT, and +P29S cells to MAPKi treatment. (**a**) For all three cell lines treated with DMSO, cells cultured on 5 kPa substrates appear to have a slight proliferative disadvantage compared to cells plated on glass. (**b, c**) Under MAPKi treatment, we note that even over the wide range of stiffness examined (5 kPa to >1GPa) that bookends the range of stiffness characteristic of traditional mechanotransduction pathways, there is minimal if any change in proliferative advantage conferred by Rac1^P29S^ expression. This contrasts the distinct stiffness-dependent proliferative advantage upon MAPKi treatment in +P29S cells observed over the stiffness range of ~200-500 Pa.

**Supplementary Figure 8.**
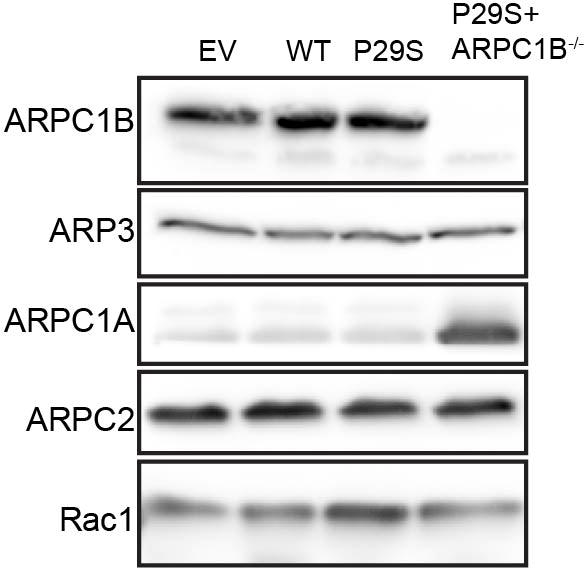
Immunoblotting to validate CRISPR/Cas9-mediated knock out of ARPC1B in +P29S cells.

**Movie S1**

Imaging of mNeonGreen-actin in A375 cells (rows: +EV, +WT, +P29S) upon MAPKi treatment (columns: DMSO, 0.003% and Dabrafenib, 33.3 nM). Images were acquired at 5-second intervals.

